# The role of standing variation in geographic convergent adaptation

**DOI:** 10.1101/009803

**Authors:** Peter L. Ralph, Graham Coop

## Abstract

The extent to which populations experiencing shared selective pressures adapt through a shared genetic response is relevant to many questions in evolutionary biology. In a number of well studied traits and species, it appears that convergent evolution within species is common. In this paper, we explore how standing, genetic variation contributes to convergent genetic responses in a geographically spread population, extending our previous work on the topic. Geographically limited dispersal slows the spread of each selected allele, hence allowing other alleles – newly arisen mutants or present as standing variation – to spread before any one comes to dominate the population. When such alleles meet, their progress is substantially slowed – if the alleles are selectively equivalent, they mix slowly, dividing the species range into a random tessellation, which can be well understood by analogy to a Poisson process model of crystallization. In this framework, we derive the geographic scale over which a typical allele is expected to dominate, the time it takes the species to adapt as a whole, and the proportion of adaptive alleles that arise from standing variation. Finally, we explore how negative pleiotropic effects of alleles before an environment change can bias the subset of alleles that contribute to the species’ adaptive response. We apply the results to the many geographically localized G6PD deficiency alleles thought to confer resistance to malaria, where the large mutational target size makes it a likely candidate for adaptation from standing variation, despite the selective cost of G6PD deficiency alleles in the absence of malaria. We find the numbers and geographic spread of these alleles matches our predictions reasonably well, consistent with the view that they arose from a combination of standing variation and new mutations since the advent of malaria. Our results suggest that much of adaptation may be geographically local even when selection pressures are homogeneous. Therefore, we argue that caution must be exercised when arguing that strongly geographically restricted alleles are necessarily the outcome of local adaptation. We close by discussing the implications of these results for ideas of species coherence and the nature of divergence between species.

## 1 Introduction

There are an increasing number of examples where different populations within a species have adapted to similar environments by means of independent genetic changes. In some cases this convergent evolution is the result of quite distinct genetic changes, involving very different genes and pathways, despite shared selection pressures; in other cases independent adaptations are identical down the same nucleotide change (Jeong & Rienzo, 2014; Stern, 2013; Martin & Orgogozo, 2013; Conte et al., 2012). Such convergent evolution within populations has been seen for many carefully studied phenotypes across a range of species, including drug resistance in pathogens, resistance to pathogens or pesticides, and the molecular basis of pigmentation changes. The phrase “parallel evolution” is also used to refer to such convergent evolution; here we use these synonymously, as we are concerned with adaptation within a single species that can occur via a different or shared genetic routes (see Arendt & Reznick, 2008, for more discussion).

The issue of convergent adaptation within species touches on a number of important questions in evolutionary biology. These include the extent to which adaptation is shaped by pleiotropic constraints (Haldane, 1932; Orr, 2005), whether adaptation is mutation-limited (Bradshaw, 1991; Karasov et al., 2010), and to what degree species should be regarded as cohesive units. Convergent evolution also affects our ability to detect adaptation from population genomic data, since no single allele sweeps to fixation over the entire area affected by the selection pressure (Pennings & Hermisson, 2006b).

Convergent evolution can occur even within a well mixed population subject to a constant selection pressure, either through selection on multiple mutations present as standing variation within the population before selection pressures switch (Orr & Betancourt, 2001; Hermisson & Pennings, 2005), or due to multiple adaptive alleles that arise after selection pressures switch (Pennings & Hermisson, 2006a). Previous work has shown that a primary determinant of the probability that multiple alleles contribute to adaptation is the product of the population size and the mutation rate (see Messer & Petrov, 2013, for a review).

Spatial population structure, as caused for example by geographically limited dispersal, also increases the chance of convergent evolution. For example, geographically patchy selection pressures can lead to much higher probability of parallel adaptation than uniform pressures, since alleles are unable to spread through intervening populations (Ralph & Coop, 2014). In Ralph & Coop (2010) we formulated a simple model of convergent adaptation in a spatially spread population with local dispersal that is exposed to some novel, spatially homogeneous selection pressure. We assumed that a single mutational change was sufficient to adapt the population after the change in environment. Under this assumption, selected alleles arise and spread locally, as shown in Figure 1. If the geographic area is large enough multiple selected alleles can arise independently and spread before any one has spread across all of space. A somewhat analogous situation also arises in spatial models of clonal inference in asexuals (Gordo & Campos, 2006; Martens & Hallatschek, 2011; Otwinowski & Krug, 2014).

**Figure 1.**
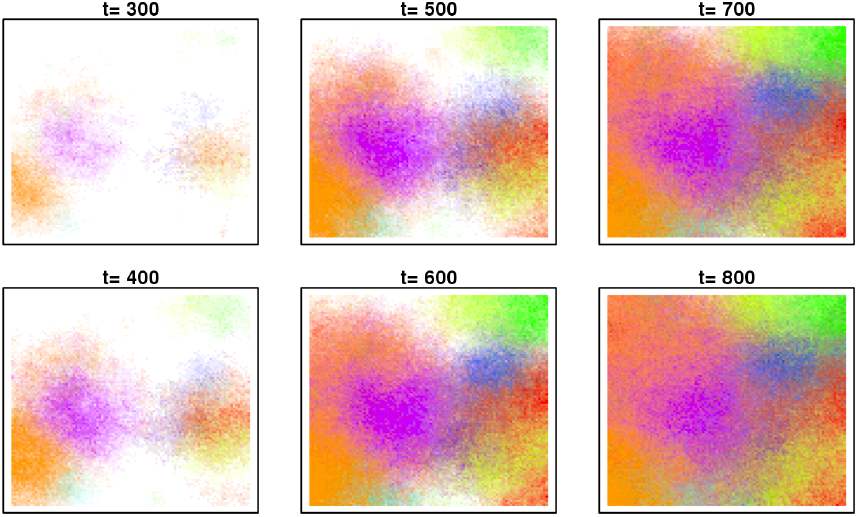
Simulation of the geographic spread of new mutations. Simulation on a grid of 101 × 101 populations with 100 haploid individuals in each, showing spread of a beneficial allele with advantage *s* = 0.1. Times are in numbers of generations, with probability of mutation per generation 0.3. Different colours show the different independent origins and spreads of selected alleles. Alleles quick establish and start to spread, but because they are selective equivalent once they spread into each other they only mix slowly (note the lack of change from generation 600 onward). More details of the simulation are given in Ralph & Coop (2010).

Under this model we previously derived the characteristic geographic scale over which multiple instances of the adaptive allele are expected to arise in parallel, a characteristic length expressed in terms the parameters of interest. In Ralph & Coop (2010) we assumed that there was no standing variation for the adaptive allele (e.g., because the allele was very strongly deleterious before the environmental change), so that parallel mutation must be due to multiple new mutations occurring after the environmental shift.

In this paper, we extend this spatial model to include standing variation present at mutation-selection balance before the selection pressures switch. Below, we show that convergent adaptation within a widespread species is likely to be common when ranges, population sizes, or mutational targets are large, as has already been seen for a number of traits. On this basis we argue that the genetics of adaptation may often be geographically local even when selection pressures are geographically broad, and that widespread selective sweeps should tend to occur only when adaptation is highly constrained (e.g. by small mutation rate or the need for a linked combination of alleles). We discuss the history, and implications, of this view for the evolutionary coherence of species and molecular evolution.

### 1.1 Model description

We assume that the species range is a large, homogeneous, one– or two–dimensional region. There are two selective classes – the *ancestral* type, and the *mutated* type (which will offer a fitness benefit after the environmental switch). We assume that separately arising mutations are distinguishable – either as selectively equivalent mutations, or by linked neutral variation. We assume that all alleles of the mutated type are selectively equivalent (both in terms how deleterious they were before, and how advantageous they are after, the environmental shift). Note that these mutations do not have to arise at the same locus, just that they are selectively equivalent: these could be mutations arising at the same base pair, or knockout mutations of any one of a number of genes in a pathway, as long as carrying at least one of these alleles is sufficient to adapt an individual to the new environment.

We also suppose that the mutated type has been at a selective disadvantage for a sufficiently long enough time in the past to be at selection-mutation equilibrium, but at a certain time the selective regime changes, so that the mutated type has a selective advantage and quickly spreads to fixation. After fixation, alleles are either descended from families of mutants present as standing variation when the selective regime changed, or from new mutants arising since that time.

For concreteness, suppose that before time *t* = 0, the mutant type has fitness 1 − *s*_*d*_ relative to the neutral type (i.e. it produces on average 1 − *s*_*d*_ times the number of offspring per generation), and that after time *t* = 0, the mutant type has fitness 1 + *s*_*b*_, where *s*_*b*_ > 0 will usually be assumed to be small, and 0 *< s_d_ <* 1. We assume that diploid fitness is additive, or at least that the important early dynamics are determined by the heterozygous fitness, with no reference to the fitness of the homozygote for the mutant alleles. Note that this means that we are ignoring the case of totally recessive alleles. We assume that the variance in offspring number is one, see Ralph & Coop (2010) for how this assumption can be relaxed. As for the other parameters, suppose that each offspring of a neutral parent is of the mutant type with probability *μ*, and that the mean squared geographic distance between parent and child is *σ*^2^. We assume that the dispersal kernel is not heavy tailed, i.e., falling into the Gaussian domain of attraction, see Ralph & Coop (2010) for discussion of heavy tailed kernels and Hallatschek & Fisher (2014) for recent progress on rapid spreading due to very long distance dispersal. The species occupies an area with mean density *ρ* of alleles per unit area (twice the number of diploid individuals per unit area).

#### Rates of origination of standing and new mutations

We make use of the commonly used approximation that neglects competition between close relatives, treating the offspring of a new mutant that appears in an area not already occupied by the mutated type as a branching process in continuous time, measured in generations (average parental age at birth). After *t* = 0, the offspring of each individual mutant thus forms (approximately) a branching process with growth rate *s*_*b*_, so each new mutant establishes locally with probability *p*_*s*_ ≈ 2*s*_*b*_. As in Ralph & Coop (2010), each new offspring has a very small probability of being a mutant and establishing locally, so the collections of times and locations at which mutants appear and establish locally is well–approximated by a Poisson process in space and time. By this we mean that there is a constant rate across space and time at which new mutations arise, and the occurrence of new mutations in any region of space-time is independent of that in non-overlapping intervals in which the mutated type has not already appeared. The rate of this Poisson process, i.e. the mean number of new mutants per unit area and per generation that appear and establish locally after *t* = 0 in areas not already occupied by the mutant type, is the product of the haploid population density, denoted *ρ*, the mutation rate (*μ*), and the probability of fixation *p*_*s*_, or approximately

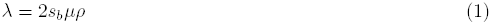

In geographic areas already occupied by the mutant type, new mutations are effectively neutral, and so unlikely to establish over the short-time scales considered here, and are hence excluded.

Before *t* = 0, on the other hand, the allele is deleterious. The genetic descendants of each new mutation are (with high probability) doomed to extinction, but may persist for a short time (note that we have assumed that *s*_*d*_ is not too small). The times and locations of new mutations before *t* = 0 also will be well– approximated by a Poisson process with rate *μρ* since mutations are rare events. Therefore, the locations of all mutant families extant at *t* = 0 whose descendants are destined to fix locally is a thinning of the original process, and so, by the Poisson Mapping Theorem, is also a Poisson process. We define *λ*_0_ to be the mean density of this process, i.e., the geographic density of standing variants that are present and escape loss when the environment shifts. If we assume that the descendants of at most only a few members of any extant mutant family at *t* = 0 will survive, and that these progenitors are near to each other in space, we can then treat each such family as equivalent to a single new mutation, but with somewhat larger probability of local establishment, following the environmental shift. (This approximation will be good if the logarithm of the size of each extant family is small relative to the establishment time, since then new families quickly “catch up” to the size of already-extant families, and the spatial distribution of each is small relative to the spread between the families.) To find *λ*_0_, consider a mutation that arose *T* generations ago, and let *Z*_*T*_ be the number of its descendants at time 0. At time *t* = 0, when the environment shifts, there are *Z*_*T*_ individuals present with the mutation, and each has probability *p*_*s*_ of establishing, approximately independently. Therefore, the probability that at least one descendant of this mutation establish and fix locally is 1 – (1 – *p*_*s*_)^*Z*^*T*, the mean number of clusters of standing variants destined to fix locally, per unit area, is

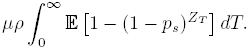

For *s*_*b*_ small, using *p*_*s*_ *≈* 2*s*_*b*_ and that 𝔼[*Z*_*T*_] = (1 − *s*_*d*_)^*T*^, we know that 𝔼[(1 − 2*s*_*b*_)^*Z*^*T*] *≈* 1 − 2*s*_*b*_𝔼[*Z*_*T*_] = 1 − 2*s*_*b*_(1 − *s*_*d*_)^*T*^, resulting in the approximation

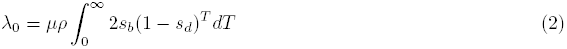

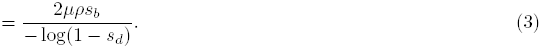

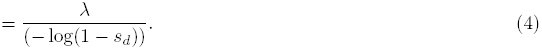

Note that for small *s*_*d*_ this can also be found by taking the expected frequency under mutation-selection balance *μ/s*_*d*_ (Haldane, 1927, 1937) multiplying it by the population density to obtain the expected number of chromosomes per unit area carrying the deleterious allele, with each of these having a probability 2*s*_*b*_ of escaping loss (an analogous approach to that taken by Orr & Betancourt, 2001).

#### Geographic spread of alleles

Once an allele has become locally established it can begin to spread across space. We assume that the allele, once established, quickly settles down to spread spatially as a traveling wave of constant speed. The behavior of this wave of advance of a beneficial allele was first described by Fisher (1937) and Kolmogorov, Petrovskii & Piscunov (1937). Under reasonably general conditions, the speed of advance of this wave is 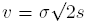. See Ralph & Coop (2010) for a more thorough review of these travelling waves. Note that the speed of the wave will vary with details of the space that individuals migrate across (e.g. see Slatkin, 1976; Slatkin & Charlesworth, 1978, for comparisons to migration on discrete grids).

#### Putting it together

Now, we can put these ingredients together for a simple model of the geographic spread of alleles, a cartoon example of which is shown in Figure 2. Initially, when the selection pressures change at *t* = 0, a set of standing variants can start to spread having escaped loss through drift. The originating mutations of these variants are depicted by lightning bolts, and occur at a density *λ*_0_ across space. They spread at velocity *v*, carving out cones in space-time. As these alleles proceed in their geographic spread, other new alleles can arise and become established in parallel, whose origins are indicated by stars. These new mutations arise and become established at rate *λ*.

**Figure 2.**
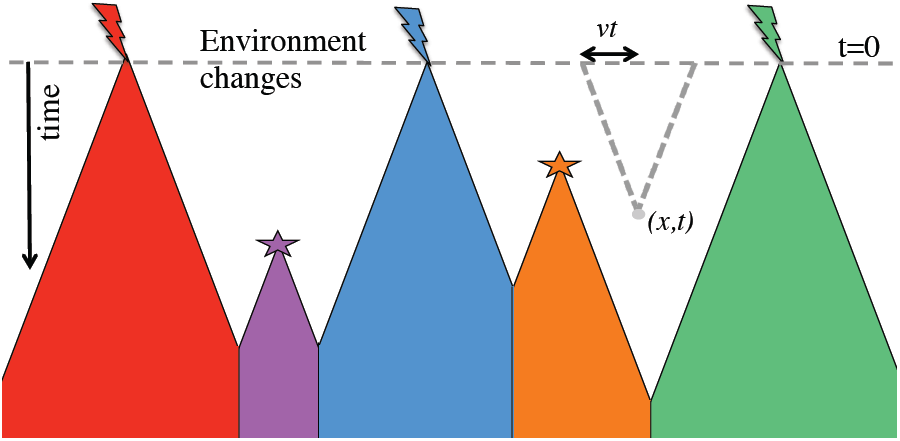
Cartoon space-time diagram of the geographic spread of standing and new mutations. Time runs down the page, and a single spatial dimension runs across the page. The environment switches at *t* = 0, following which adaptive alleles that successfully escape loss by drift spread locally from both standing variants (lightning bolts) and new mutations (stars). (Note that the standing variants arose at times prior to *t* = 0.) The mean spatial density of successful standing variants at *t* = 0 is *λ*_0_, and the mean density in (space × time) of successful new mutations is *λ*. These successful alleles spread spatially outwards at speed *v*, so the space-time profile of their spread forms a cone. Any mutation that arises in an area already within a cone will not have a selective advantage. When these adaptive alleles meet, their rapid advance is halted because they no longer have an advantage, so their boundaries form straight lines on short time-scales (over longer time-scales, they will mix into each other). One of the ways we derive various results in the text is by taking a space-time point (*x, t*) and asking the probability that no successful alleles have spread there yet. The grey dashed line shows a cone radiating backward in space-time, originating at (*x, t*), with slope *v*; its radius at *t* = 0 is *vt*. This shows the location *x* is not yet adapted at time *t* because no successful alleles appear within this grey cone.

As we outlined in Ralph & Coop (2010), this model of geographic convergent evolution, when *λ*_0_ = 0, is analogous to a model of crystallization due to Kolmogorov (1937). In this model, nucleation sites form at random at a constant rate in time and space and initiate the radial growth of new crystals. After their initial spread, the different orientations of crystals form a random tessellation of space, whose properties have been studied by Møller (1992, 1995) and others (Bollobás & Riordan, 2008; Gilbert, 1962). The generalized version of this process, for non-constant wave speeds and inhomogeneous Poisson processes is known as the Kolmogorov–Johnson–Mehl–Avrami tessellation (Fanfoni & Tomellini, 1998). Our combined process with both standing variation and new mutation is a special case of the KJMA tessellation, where the spatial-temporally homogeneous Poisson origination process of new mutations, is supplemented by a single pulse of origination points at time zero with spatial density *λ*_0_. (For the purposes of analogy, we could imagine that before time *t* = 0 the temperature is high enough that nucleation sites appear but do not persist long.)

If we ignore the effects of new mutation, then everything about the process is relatively simple: each point in space will be first reached by the wave whose origination point lies closest to it. This random tessellation of space is known as a Poisson-Voronoi tessellation (Møller, 1994) (i.e. the cells formed by assigning regions of space to the nearest point in a Poisson process). The properties of this tessellation by alleles is determined by the spatial locations of the initiation points, which are sampled from a spatially homogeneous Poisson process, independent of the rate of spatial spread. Introducing new mutations cause some qualitative changes beyond dependence on new parameters: the cells formed by a Voronoi tessellation have straight sides, but the introduction of new mutations cause these to curve (because the radii of the colliding circles differ; see Figure 1 of Ralph & Coop, 2010, for a graphical depiction of this point).

### 1.2 G6PD example

Below we describe a number of properties of our process, but we will first introduce a motivating example to provide some concrete numbers to use for illustrative purposes.

Over roughly the past ten thousand years, alleles conferring resistance to malaria have arisen in a number of genes and spread through human populations in areas where malaria has been endemic (Kwiatkowski, 2005). A number of these alleles appear to be examples of convergent adaptation, as different derived mutations in the same gene are seen in different individuals. For example, a number of changes that confer malaria resistance have been observed in the *β*-globin gene; and the sickle cell allele may plausibly have arisen by up to five independent occurrences of the same base pair mutation at different locations within Africa (Flint et al., 1998; Ralph & Coop, 2010). Another particularly impressive case of convergent evolution is presented by the numerous changes throughout the X-linked G6PD gene, with upward of 50 polymorphic variants (above 1% local frequency) having so far been described that lower the activity of the enzyme (Howes et al., 2013; Minucci et al., 2012). These alleles are now found at a combined frequency of around 8% frequency in malaria endemic areas, rarely exceeding 20% (Howes et al., 2012). Whether these all confer resistance to malaria is unknown, but malaria is thought to be the primary driver of these polymorphisms (see Hedrick, 2011, for a general review). Three G6PD deficiency alleles are particularly common and relatively well studied: the *A*– allele found in much of sub-Saharan Africa; the Med allele found in the Mediterranean and Middle East; and the Mahidol allele found Myanmar and Thailand. The protective effects of these G6PD alleles are complicated, and are likely heterogeneous across study populations and the form of malaria considered (see Manjurano et al., 2015; Malaria Genomic Epidemiology Network, 2014, for recent discussion). The *A*– and Mahidol alleles are thought to offer some protective effects against *Plasmodium falciparum* and *P. vivax* in both heterozygote females and/or hemizygote males (Ruwende et al., 1995; Louicharoen et al., 2009; Manjurano et al., 2015). Haplotype-based analysis of genetic diversity surrounding *A*–, Med, and Mahidol suggest that they have spread over the past few thousand years (Tishkoff et al., 2001; Slatkin, 2008; Saunders et al., 2005; Louicharoen et al., 2009), consistent with the age of other known malaria resistance alleles. Population genetic analyses suggest that these three variants each have a hemizygote/heterozygote selection coefficient of 0.05 − 0.3 (Tishkoff et al., 2001; Slatkin, 2008; Saunders et al., 2005; Louicharoen et al., 2009). This is in reasonable agreement with the selection coefficients calculated by Ruwende et al. (1995) on the basis of the present day levels of resistance to malaria due to the *A*– allele.

Given such strong selection the alleles should have risen quickly to fixation, so their presence at intermediate frequency, over a broad geographic area, makes it a good candidate for a recently balanced polymorphism due to heterozygote advantage (note that the conditions for a balanced polymorphism are complicated by the hemizygosity of males, see Hedrick, 2011; Pamillo, 1979). Indeed, hemizygous males and homozygous females suffer from G6PD deficiency, and homozygote females may also not be protected against malaria (Manjurano et al., 2015; Malaria Genomic Epidemiology Network, 2014). The theory we use regarding the “wave of advance” (Fisher, 1937) applies as well in the case of heterozygote advantage (Aronson & Weinberger, 1975), with the selected allele spreading locally to the equilibrium frequency (rather than fixation). Therefore, our framework is applicable to the spread of G6PD, with speed determined by the advantage of heterozygotes when rare. We assume that before malaria became prevalent, G6PD deficiency alleles suffered a decrease in relative fitness of *s*_*d*_ in heterozygote and homozygotes females and hemizygote males. Assuming that the underlying causes and strength of this drop in fitness have not changed, we estimate that *s*_*d*_ has to have been upward of ∼ 0.05 (if *s*_*b*_ ≥ 0.05), in order to have resulted in the equilibrium frequency seen today in areas with endemic malaria (based on heterozygote advantage calculations for the X chromosome, results not shown, see also Ruwende et al., 1995).

The geographic area of Central and Eastern Asia with malaria is on the order of ten million square kilometers. In that area there are at least 15 common, clinically relevant variants (see Figure 3, from Howes et al., 2013). (These are type 2 variants that express at < 50% enzyme activity, predispose individuals to haemolytic anaemia, and are found in at least 10 localities; see Howes et al. (2013) for more details.) Therefore, the average width of an area occupied by an allele is 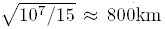. The coding region of G6PD is 515 codons long, and around 140 distinct deficiency alleles have been observed. Assuming a mutation rate of ≈ 10^−8^ per base pair per generation, we can take as an order-of-magnitude estimate *μ* ≈ 10^−6^ per generation. The dispersal and demographic parameters of humans in the past few thousand years is unclear, particularly as we are concerned with the “effective” population density (i.e. population density divided by variance in offspring number). We therefore will use two reasonable values for the effective population density: *ρ* = 2 and 0.2 people per km^2^, and three values for the dispersal distance: *σ* = 10, 50 and 100 kilometers per generation. Clearly, human migration has been shaped both by local dispersal and larger-scale expansions (see Pickrell & Reich, 2014, for a recent discussion), so these parameters only provide a rough view of the process.

**Figure 3.**
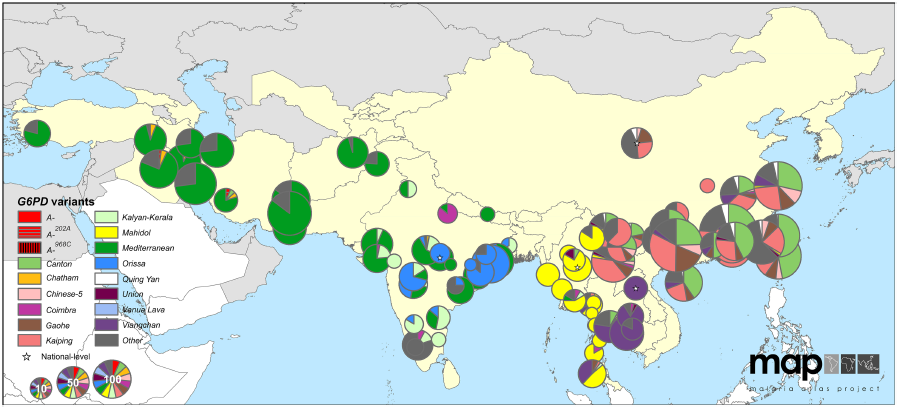
Map of G6PD-deficiency allele frequencies across Asia. The pie chart shows the frequency of G6PD-deficiency alleles. The size of the pie chart indicates the number of G6PD-deficient individuals sampled. Countries with endemic malaria are colored yellow. Figure taken from Howes et al. (2013) http://www.malariajournal.com/content/12/1/418.

### 1.3 The geographic resolution of adaptation from new and standing variation

In Ralph & Coop (2010), studying the model without standing variation, we defined a *characteristic length* which gave the spatial scale across which mutants with distinct origins would establish. This was proportional to the mean distance between neighboring established mutants, but had the advantage of being easier to calculate. Furthermore, the time scale over which adaptation occurred could be found by dividing the characteristic length by the speed at which the mutants spread. We first define a similar characteristic length for this new model.

Suppose we fix our attention on a particular new mutation that happens to be the first to occur in some region. If it does not encounter other locally fixed, beneficial mutants, it will cover a distance *L* in time *L/v*. In doing so (in two-dimensions) it will have started from a point in space and spread out to cover a circular geographic region of area *πL*^2^. The cylinder in space-time with this circle as a base and height *L/v* has volume of this cylinder is *πL*^2^ × (*L/v*). Therefore, the number of other successfully established mutations that would have appeared in the circle it has covered up until this time is Poisson with mean *λ*_0_*πL*^2^ + *λπL*^3^*/v* in two dimensions (and 2*λ*_0_*L* + 2*λL*^2^*/v* in one dimension). Therefore, if we define *χ* for a two-dimensional model to be the unique positive solution to

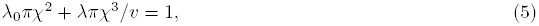

then *χ* gives the distance spread unobstructed by the descendants of a new mutant before it is expected that one other successful mutation would have arisen in the area covered so far. The explicit formula for *χ* in two dimensions can be found by rearranging eqn. (5) but is cumbersome; here we omit it. In one dimension things are a little prettier and the characteristic length is 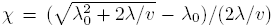. Substituting expressions for *λ*_0_, *λ*, and *v* from above, in two-dimensions we can rewrite eqn. (5) as

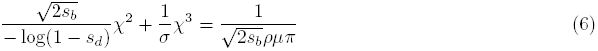

From this we see that *χ* decreases with *ρ* and *μ*. Furthermore, for large *σ*, the characteristic length approaches the value we would obtain just from standing variation:

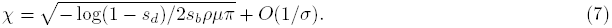

On the other hand, if the mutant allele is highly deleterious before *t* = 0, then standing variation is unimportant the characteristic length is approaches the value from Ralph & Coop (2010):

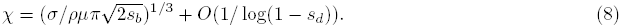

These two end points help build our intuition for the interaction of parameters in shaping the geographic scale of convergent evolution. By the above calculation, we know that the relevant mutations occur about distance *χ* apart, and occur within the first *χ/v* generations. Said another way, if we look in a circular region of space of radius *χ* over *χ/v* generations, we expect to find roughly one mutational origin.

In Figures 4 and 5, we show the range of characteristic lengths as a function of various parameters chosen to match the evolution of malaria resistance at G6PD. These curves match our intuition that higher population densities result in smaller characteristic lengths (as would higher mutation rates). Allowing standing variation increases the input of new alleles and so decreases the characteristic length below that predicted by only new mutations (i.e. equation (8), Ralph & Coop (2010)). In turn, increasing the prior deleterious effect of the allele (*s*_*d*_) acts to increase the characteristic length until it reaches that predicted by new mutation alone. Higher dispersal distances lead to larger characteristic lengths, since more rapid geographic spread block other mutations from establishing. A larger selective advantage (*s*_*b*_) acts in two conflicting ways: aiding the rapid geographic spread of established alleles, and also helping more independent copies to escape drift and become established. The effect of helpinglocally establish alleles wins out, since increasing the selective benefit *s*_*b*_ decreases the characteristic length. (This can be shown in general by differentiating (6) with respect to 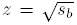, showing that *∂_z_χ* = –(*C*_1_*zχ* + *C*_2_*χ*^2^)/(*C*_3_*z*^2^ + *C*_4_*zχ*) ≤ 0 for appropriate nonnegative constants *C*_1−4_.) This effect is strongest when only standing variation contributes (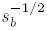, equation (7)), as in that case the speed of spread does not matter only the initial density of established alleles after the environmental shift. The dependence of the characteristic length on *s*_*b*_ when only new mutations contribute is much weaker (of order 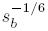, equation (8)) Overall, the range of characteristic lengths observed are reasonably consistent with the average diameter of a G6PD variant in Eurasia of 800km, especially for the lower population density, as long as the fitness cost of G6PD-deficiency alleles before malaria (*s*_*d*_) was high.

**Figure 4.**
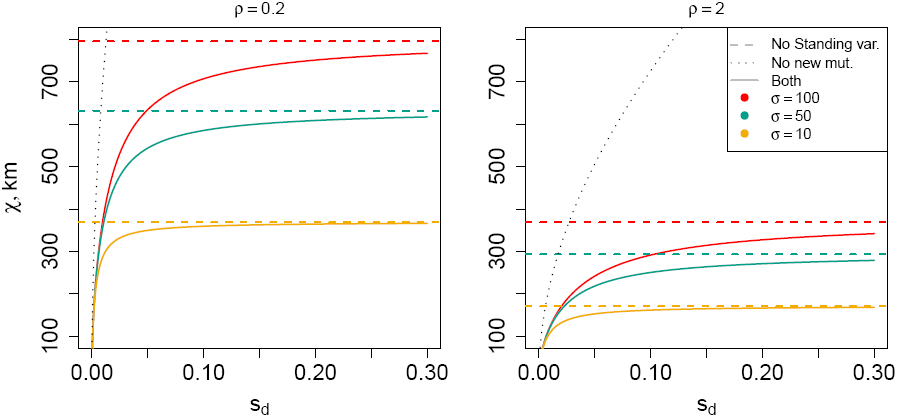
**Characteristic length, in kilometers, as a function of selective disadvantage**, compared to the corresponding quantity without standing variation (equation (8)) and to the quantity only considering standing variants (equation (7)). The other parameters are chosen to match those of G6PD, *μ* = 10^−6^ and *s*_*b*_ = 0.05. As the characteristic length with only standing variation is independent of dispersal distance it is plotted as a single black dotted line.

**Figure 5.**
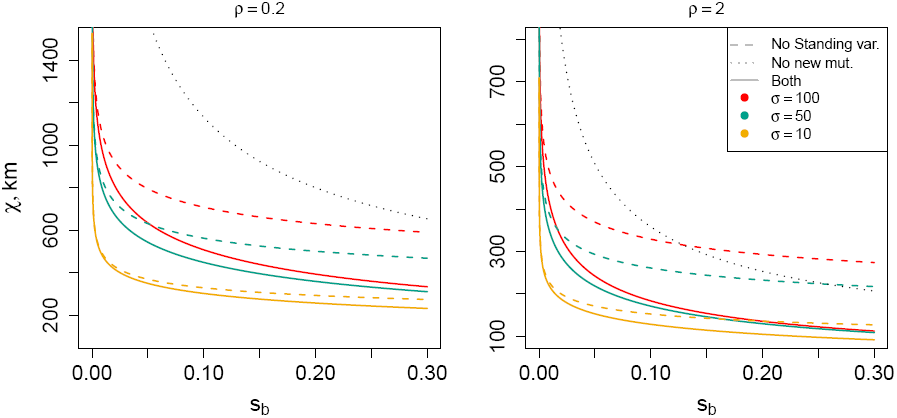
**Characteristic length**, in kilometers, as a function of selective advantage, at two population densities, holding the prior disadvantage of the allele fixed at *s*_*d*_ = 0.05 and the mutation rate fixed at *μ* = 10^−6^. We compare this to the corresponding quantity without standing variation (equation (8)) and to the quantity only considering standing variants (equation (7)).

Finally, while the form of eqn. (6), specifying the characteristic length, is not particularly intuitive we can use it to ask when we should expect multiple adaptative alleles within a large geographic region where our selective pressure is present (thanks to Sam Yeaman for proposing this interpretation). Consider the case where this geographic area is a fairly regular shape of area *G* and diameter 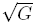. Denote the total effective population size over this area by *N* = *ρG*, and the standard deviation of dispersal as a fraction of the diameter of this area by 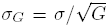. Measuring distance in units of the diameter of 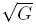, we expect multiple mutations when *χ* < 1 a condition which will be met when

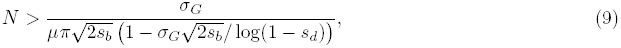

this is found by rearranging eqn. (6) having set *<* = 1. This nicely shows that *σ_G_/μ* is a primary determinant of the critical population density necessary for convergent evolution, when *σ_G_* is small. As *σ_G_s_b_/s_d_* becomes large, e.g. because the allele is not too deleterious before the environmental shift, we get our standing variation only case where the dependence of *σ*_*G*_ drops out leaving the critical population size as 1*/*(2*μπs_b_/s_d_*)

Note that the conflicting roles of *ρ* and *σ* mean that even in species where levels of neutral differentiation are low, geographic convergent adaptation may be common. This is because low levels of neutral genetic differentiation between geographic regions can be due to high population densities rather than high dispersal distances, and high population densities would allow convergent adaptation. As such geographic convergent evolution may be common even in species with little neutral population structure.

### 1.4 Time to adaptation

It is also straightforward to compute the mean time until adaptation. Imagine a geographic location, and let *τ* ≥ 0 be the time at which this location is first reached by some advantageous mutation. Then, as can be seen from the perspective of the grey dot in Figure 2, *τ > t* if and only if the cone with point at (*x, t*) and slope *v* extending back to *t* = 0 (light grey dashed lines) is empty of successful mutations. Since we assume that successful alleles arise as a Poisson process, in two dimensions

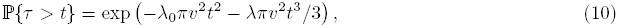

i.e., the combined probability that area of a circle of radius *vt* surrounding our point at *t* = 0 was free of successful standing variants, and no successful, new mutations arose in the cone (that has radius *r* = *vt* at *t* = 0 and height *h* = *t*, and hence volume *πr*^2^*h*/3). Since 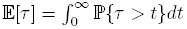,

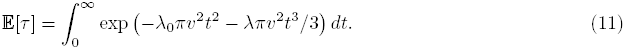

For applications we evaluate this integral numerically.

In Figure 6 we show the mean time until adaptation for various values of the parameters chosen to match the case of adaptation at G6PD. Increasing *σ* and decreasing *s*_*d*_ lower the time to adaptation, as alleles spread geographically more quickly and are present as standing variation more often respectively. Increasing *s*_*b*_ strongly decreases the time to adaptation, as it both causes more alleles to escape drift and to rapidly spread. Given that the G6PD alleles likely spread over a few thousand years, i.e. less than a few hundred generations, this time scale seems quite plausible, except perhaps for the lowest dispersal distances.

**Figure 6.**
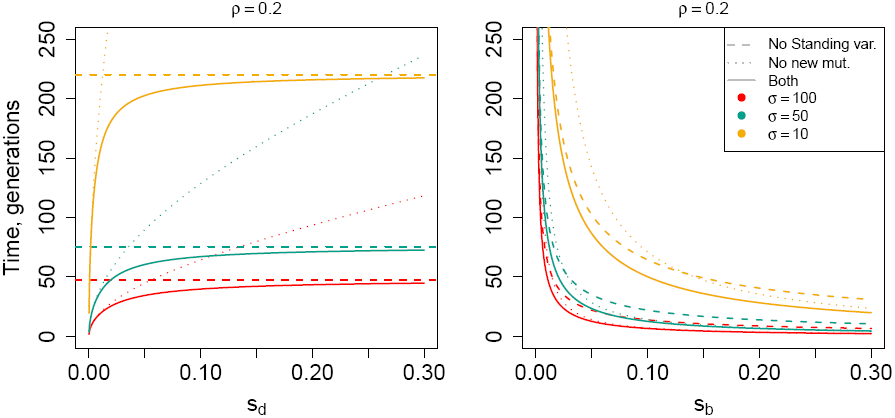
**Mean adaptation times** as a function of selective advantage and disadvantage. We compare these to the corresponding quantities without standing variation and only considering standing variants.

### 1.5 The contribution of standing variation.

We can also ask in our framework to address what proportion of new adaptive variants arise from standing variation. We have defined *λ*_0_ to be the mean density of standing variants that are present and escape loss when the environment shifts. We will define *γ* to be the mean density of newly arising alleles that spread having arisen in an area free of other adaptive alleles. Since the probability that a mutant arising at location *x* and time *t* is lucky enough to be born in a location not already occupied by mutants is ℙ{*τ > t*}, we can see 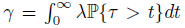, and hence *γ* = *λ*𝔼[*τ*]. Therefore, the mean proportion of adapted patches that come from standing variation is

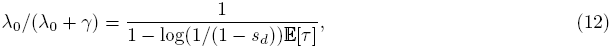

using the fact that *λ*_0_ = *λ/* log(1*/*(1 – *s*_*d*_)). There are *λ*_0_ + *γ* patches per unit area, so the typical patch (informally, the patch around a randomly chosen successful mutation; formally, drawn from the Palm measure (Cox & Isham, 1980)) occupies area 1*/*(*λ*_0_ + *γ*).

We can also find the mean proportion of space covered by standing variants (we will restrict ourselves to two dimensions). At time *t* a geographic location has not yet been reached by the mutation with probability given by equation (10). Given that it has not been reached by *t*, the probability that it will be reached by time *t* + *dt* by a standing variant is approximately 2*λ*_0_*πv*^2^*tdt*, which is *λ*_0_ multiplied by the thin slice of extra area in our expanded circle at *t* = 0, which has gone from a radius *vt* to *v*(*t* + *dt*). The corresponding probability that the point is reached by a new variant is *λπv*^2^*t*^2^*dt*, which is *λ* multiplied by the sliver of extra volume in our space-time cone at time *t* + *dt* compared to that at time *t*. Therefore, the mean proportion of space covered by standing variants, is

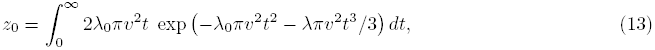

this is the probability a given location is reached first by a standing variant (which follows from competing our two exponential waiting times). For applications we evaluate this integral numerically.

Furthermore, if we define *a*_0_ to be the mean area occupied by a typical standing variant, then *a*_0_ is given by the proportion of the range occupied by standing variants divided by the mean density of unique standing variants, i.e. *a*_0_ = *z*_0_*/λ*_0_. We can solve for *a*_+_, the corresponding mean area occupied by a given new variant, using the formula *a*_0_*/a*_+_ = *z*_0_*/*(1 − *z*_0_).

In Figure 7 we show the proportion of alleles that spread from standing variation, and the proportion of geographic space covered by standing variants for parameters chosen to match our G6PD example. Even for relatively large deleterious costs prior to the environmental switch, standing variants still make up quite a large proportion of the adaptive alleles, and an even larger proportion of the range (they occupy a larger area than new mutations, since they get a head start).

**Figure 7.**
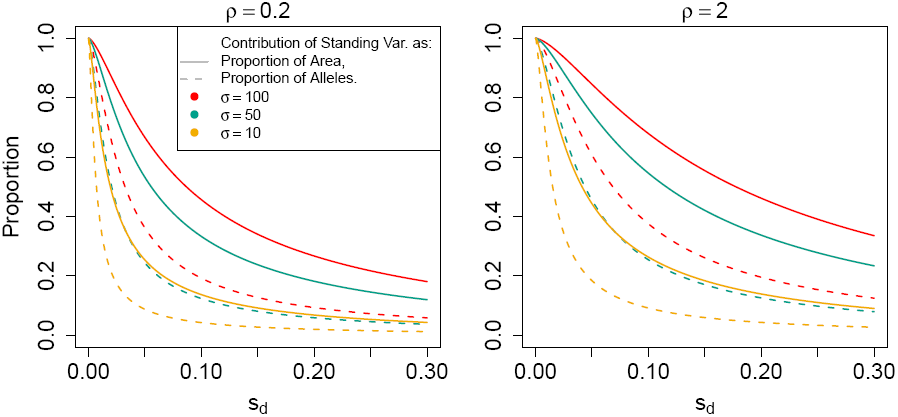
**Mean proportion of patches arising from standing variation**, as a function of selective disadvantage, at two population densities. The parameters given are for the G6PD example. Both panels show the expected proportion of patches that arise from standing variation (dotted lines, equation (12)), and the expected proportion of geographic space that is covered by adaptation from standing variation (solid lines, equation (13)). We set *μ* = 10^−5^ and *s*_*b*_ = 0.05 to match our G6PD example.

### 1.6 Multiple variant types

Another problem that we can address with this work is the extent to which pleiotropy biases adaptation towards the repeated use of particular subset of loci (i.e. convergence at genetic level). While many alleles may confer the beneficial phenotype, not all will contribute equally to adaptation if they have negative pleiotropic consequences. There are at least two ways that negative pleiotropy can contribute to high rates of convergence when adapting if a single change is sufficient for adaptation. First, negative pleiotropic effects can reduce the overall beneficial selection coefficient of an allele in the new environment, making them unlikely to become established and slow to spread (and in the worst case making them deleterious). This first effect has been well studied by a number of authors (Orr, 2000; Otto, 2004; Welch & Waxman, 2003; Chevin et al., 2010) and its role in genetic convergence examined (Orr, 2005; Chevin et al., 2010; Unckless & Orr, 2009). A second contribution is that alleles that have less negative pleiotropy are more likely to be present as standing variation before the environmental shift, and so are more able to respond immediately.

Here we focus primarily on the second effect. Let’s imagine for the moment that there are several classes of beneficial allele, all having the same beneficial selection coefficient (or at least that beneficial selection coefficients are similar enough that our selective exclusion approximation holds over the time scale on which we examine the process). Each class of mutations *j* has its own mutation rate *μ_j_* and selective disadvantage *s*_*d,j*_ prior to the environmental switch. As they have the same beneficial selection coefficient after the switch, all of the waves travel outward at a rate *v*. Then, the density of type *j* standing variants per unit area and the input rate of *de novo* variants per unit area per generation are, respectively,

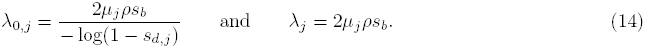

Using these rates, and an argument analogous to that used to derive equation (13), at the time when every location has been reached by an adaptive allele, the proportion of the species range covered by alleles of type *j* is

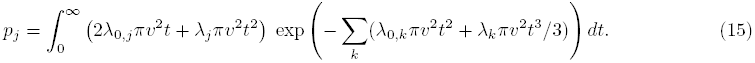

If we only allow standing variation this collapses to

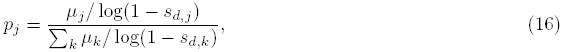

while if we only allow new variation, i.e. if all variants are highly deleterious before the environment switches, *p*_*j*_ = *μ*_*j*_/(∑_*k*_ *μ*_*k*_).

To illustrate some of the properties of this model, let’s imagine the somewhat extreme scenario in which there is a single base pair at which a possible mutation is relatively free of negative pleiotropy (call this class 1); and a larger mutational target where changes have more serious pleiotropic consequences in the ancestral environment (class 2). We set *s*_*d,1*_ ≤ = s_*d,2*_ = 0.05 and *μ*_1_ = 1 *×* 10^−8^, assume that both classes share a beneficial selection coefficient of *s*_*b*_ = 0.05, and think of the second class of alleles as arising at one of ten, one hundred, or one thousand base pairs. We show the expected proportion of space covered by the rarer, class 1 mutations in Figure 8. As expected intuitively, the contribution of the rarer mutation decreases as the mutational target of the second class becomes larger, and as the difference in the negative pleiotropic consequences of the two classes of alleles decreases. The case with standing variation only is the best case scenario for the rarer mutation, so its rate of introduction after *t* = 0 is necessarily lower. However, the standing-variation-only case does seem to provide a reasonable rule of thumb, especially for parameter combinations, such as high population densities and high dispersal distances, that increase the contribution of standing variation (and similarly for high *s*_*b*_).

It is natural to also incorporate differences in the beneficial selection coefficients of the different classes of alleles, to allow for negative pleiotropic effects acting to suppress the advantage of an allele once the environment switches. One simple way to do this is to simply replace *s*_*b*_ with *s*_*b,j*_ resulting in a class-specific new allele establishment rates (*λ*_*j*_) and rates of spread (*v*_*j*_). These then could be used in equation (15). For instance, we could extend the two class model above so that class one has the additional advantage of *s*_1*,b*_ *> s*_2*,b*_. This would further increase the contribution of the rarer class, because this class would both overcome drift more often and spread more rapidly. However, a straightforward application of the logic of used to construct equation (15) fails once the different allelic types meet, since the assumption of selective exclusion no longer holds: alleles with higher *s*_*b*_ will spread, at a lower speed, into regions occupied by alleles with lower *s*_*b*_. Given enough time, the most advantageous type (type 1, in this case) would spread everywhere, and so substituting multiple values for *s*_*b*_ in equation (15) would only provide a short-term approximation to a longer term dynamic. Even if the initial tessellation has formed with purely class 2 alleles, the first allele would*v*have a selective advantage *δs* = *s*_*b,1*_ – *s*_*b,2*_, and so would arise at rate 2*ρμ*_1_*δs* and would spread at speed 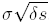. An extension of our Poisson process model could incorporate these effects, by thinning the Poisson process of establishing mutations by correctly, but is considerably less tractable. Whether allele 2 persists would depend on the linkage arrangements between loci. If the loci underlying allele 1 and 2 are unlinked, then allele 1 can spread without disrupting allele 2. However, if they are linked, the spread of allele 1 may push allele 2 out of the population. More complicated dynamics, including spatial Dobzhansky-Muller incompatibilities (Kondrashov, 2003; Ralph & Coop, 2010) could ensue if there are epistatic interactions between the alleles.

### 1.7 Local establishment and comparison to panmixia

In the above, we have assumed that once the mutation appears, conditional on eventual fixation, it begins to spread spatially at speed *v* instantly, effectively neglecting the time it must first spend escaping demographic stochasticity. In Ralph & Coop (2010) we addressed this by noting that there would be no change at all in our results if all mutations had to wait the same amount of time before fixing locally, and that this time was short relative to the time it took the wave to spread across the characteristic length; we then showed via simulation that this was reasonable in certain situations. In this section we examine this assumption in more detail, although mostly through heuristic arguments, and also compare the results above to the results without geographic structure of Pennings & Hermisson (2006a).

We are assuming that shortly after a new mutation appears, it can be approximated by a branching process growing at rate *s*_*b*_ until the point that it grows large enough to “feel” spatial structure, at which point it begins to spread as a more–or–less deterministic wave. Although we are not aware of good analysis of this transition, the relevant size of the branching process when spatial structure becomes important should be something close to *σ*^2^*ρ* (i.e. Wright’s “local effective population size”, Wright (1943)). Let *Z*_*t*_ be a continuous-time branching process with *Z*_0_ = 1 and 𝔼[*Z*_*t*_] = *e*^*s*^_*b*_*t*. Then we know that there exists a random variable *W* such that lim_*t→∞*_ *e*^*−s*^_*b*_^*t*^*Z*_*t*_ = *W* almost surely, so that if *τ* is the time *Z*_*t*_ reaches size *σ*^2^*ρ*, then *σ*^2^*ρ* = *Z*_*τ*_ *≈ e*^*s*^_*b*_^*τ*^ *W* (Jagers, 1975). From this we know that *τ* ≈ (1*/s_b_*)(log(*σ*^2^*ρ*) – log *W*); although more detailed information is available (e.g. a central limit theorem for *τ*, Nagaev (1971)), we will stick to the loose interpretation.

So, roughly speaking, we need to evaluate the importance of a delay of about *T* = (1*/s_b_*) log(*σ*^2^*ρ*). New mutations will appear and become established during this time if 2*ρσ*^2^*μs_b_* ≥ 1*/T*, i.e. if 2*ρσ*^2^*μ* ≥ 1*/* log(*σ*^2^*ρ*). Our model will still be a good approximation, however, as long as *T* is short relative to the time a wave takes to spread between nearby mutational origins. This can be worked out, but it is simpler to note that if the converse is true (i.e., that reaching local fixation is slow compared to the spread of the wave), the process is largely unaffected by spatial structure, and so the panmictic model is a good approximation for the true process.

Pennings & Hermisson (2006a) show that under a panmictic model with certain assumptions on the parameters, the number of independent origins due to both standing variation and new mutation seen in a sample of size *n* has approximately the Ewens distribution with parameters *n* and *θ* = 4*N μ*. As *n* increases, the total number of types seen grows as log *n*. In our model, we can increase total population size by increasing either the density *ρ* or the total amount of area. In either case, the predicted number of distinct types grows linearly with *n*, much faster than under panmixia.

The results of Hermisson & Pennings (2005) and Pennings & Hermisson (2006a) suggest that in a panmictic population the number of independent alleles (and their frequencies) in a sample is nearly independent of *s*_*b*_ and *s*_*d*_ (although this breaks down with fluctuating population size, Wilson et al., 2014). In the panmictic model the lack of dependence on *s*_*b*_ comes about because while increasing *s*_*b*_ increases the rate at which independent mutations become established, it also accelerates the frequency gain of established alleles, hence decreasing the time period in which new alleles can arise and hope to be at significant frequency in the population. These two effects approximately cancel each other out leading to no strong effect of *s*_*b*_ on the number of independent alleles. Decreasing *s*_*d*_ increases the number of standing variants within a population, increasing the number of alleles that manage to establish and spread from standing variation (Hermisson & Pennings, 2005; Orr & Betancourt, 2001). However, having more established standing alleles acts to exclude the spread of new alleles that arise once the environment switches. These two opposing effects again cancel out, leading to little overall effect of *s*_*d*_ on the number of independent alleles. In contrast, our results show that the characteristic length (closely related to the density of independent alleles) depends on both *s*_*b*_ and *s*_*d*_ in a geographically spread population. Like the panmictic model, in our model, alleles also act to exclude each other; however, the geographic spread of an allele is slow compared to the initial exponential growth of an allele in a panmictic population. That means that the role of selection in helping alleles become established can dominate, leading to more independent origins, both by being weaker before and stronger after the environmental switch.

## 2 Discussion

When the geographical area where a species experiences a selection pressure is greater than our characteristic length we expect multiple independent alleles to arise and spread in that area. Our results suggest that convergent evolution among populations may be quite common, at least when population densities and mutational target sizes are not too small, and dispersal is limited across the range. While considering smaller mutational target sizes (e.g., less than the 100bp in our example) or lower population densities would lower the probability of convergence within a given geographical area, it would also act to considerably lengthen the time to adaptation. In such cases adaptation may simply fail to occur at all or populations may adapt via some other means (e.g., by using a broader mutational target).

Our inclusion of standing variation before environmental change greatly increase the probability of convergence within a species. While at face value this increase seems unsurprising, this relationship differs markedly from the case of multiple competing alleles in a panmictic population (see discussion above and in Hermisson & Pennings, 2005; Pennings & Hermisson, 2006a). Importantly, allowing standing variation may greatly lower the time until the species becomes adapted across the geographic range of the selection pressure. We have also shown that adaptation through standing variation biases the type of variation towards those alleles with fewer pleiotropic effects, since these are more common as standing variation before the environmental shift. This bias can in some cases easily overcome quite significant differences in mutational target sizes among loci allowing the same locus to be repeatedly the source of adaptation even if there are seemingly many different routes to adaptation. (See Figure 8.)

**Figure 8.**
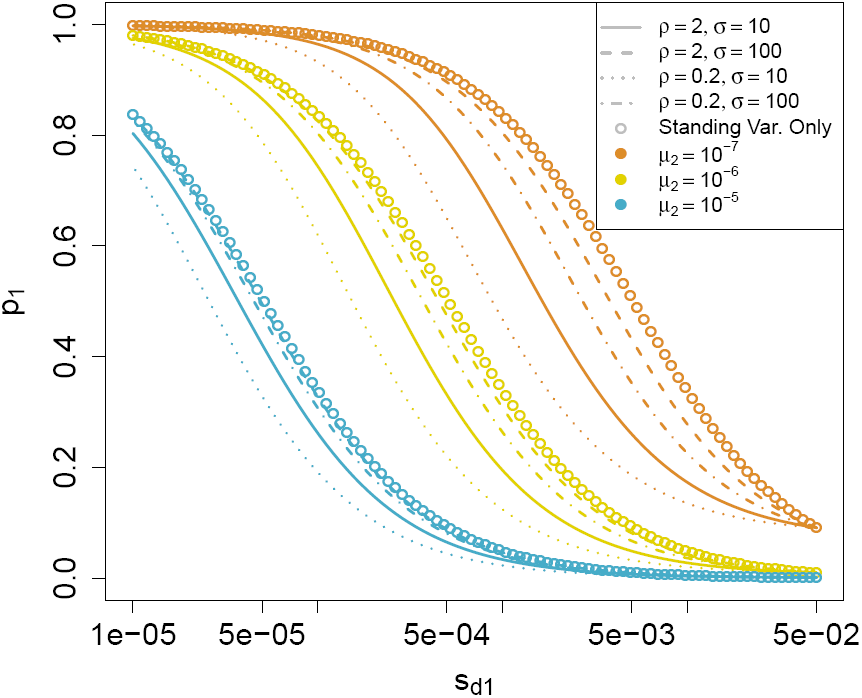
Proportion of space covered by rarer but less negatively pleiotropic mutation. Empty circles give the result for standing variation only, equation (16). Lines give the result allowing both standing and *de novo* mutations (equation (15)). Here we hold *s*_*b*_ = *s*_*d,2*_ = 0.05 and *μ*_1_ = 1 *×* 10^−8^.

### The confusing signal of geographic convergent evolution

As we have argued in Ralph & Coop (2010) the ease with which geographic convergent adaptation occurs means that we should incorporate it more widely into our thinking about the genetic basis of adaptation. For example, the absence of European skin pigmentation alleles in ancient DNA from Europeans who lived several thousand years ago has led to the suggestion that these individuals had dark pigmentation (Olalde et al., 2014; Lazaridis et al., 2014; Wilde et al., 2014). However, given our results and the partially convergent basis of skin pigmentation between Europeans and East Asians (Norton et al., 2007; Edwards et al., 2010) it seems just as plausible that these ancient individuals adapted to high latitudes via a different complement of “light-skin” pigmentation alleles; to our knowledge, we have no strong evidence either way. Such convergence may considerably complicate the exploration of phenotypes and adaptation among populations using variants mapped in a limited set of populations (Berg & Coop, 2014).

More generally, if geographic convergence is common, we should often expect to see selected alleles that are strongly geographically restricted as they have simply not had time for neutral gene flow to spread them across the landscape. Such convergent alleles will be *F*_*ST*_-outliers compared to older neutral variation, until there is sufficient time for migration to smooth out them out across the landscape. This pattern may be very hard to distinguish from local adaptation using population genomic approaches alone. This is especially problematic as boundaries between convergent alleles may often occur where gene flow rates are low, i.e. historical and ecological breaks, even if the alleles concerned have no bearing on the ecological differences across these breaks (see Bierne et al., 2011, for a wide-ranging discussion of how allelic differentiation may build along particular zones). We are rarely so fortunate as to know as much about the genetics, phenotypic distributions, and potential selection agents as we do for malaria resistance in humans. Therefore, we must be wary of mistaking the strange spatial distributions of particular alleles for adaptation to some very specific selection pressure (e.g. the distribution of the Mahidol allele in Figure 3), when they are simply elements of a larger geographic mosaic of alleles responding to a broadly shared selection pressure.

Each of these local sweeps will be associated with the haplotype on which the particular allele arose. Under the parameter regime we study, standing variants are still quite young, so we do not expect a strongly reduced hitchhiking effect. As such, following the initial period of adaptation, we should expect the population to be partitioned into a set of geographically restricted long haplotypes. Given sufficient time these haplotypes will mix together through migration and drift, potentially leading to a within population signal of a sweep from multiple independent mutations if our selected allele occurs at the same locus (Pennings & Hermisson, 2006b), or to multiple partial sweeps if the loci are scattered across the genome (Coop & Ralph, 2012).

Our results are predicated on the idea that adaptive variants are initially rare within populations, i.e. they are reasonably deleterious before the environment switches. In contrast, if they adapt via common variation even distant populations could have a shared basis of adaptation, e.g. previously neutral (or nearly neutral) variation, shared among populations. If many loci contribute to variation in a trait, then selection on any one allele may be weak, which might lead adaptation to use the same alleles in different populations. However, sufficiently differentiated populations may still adapt via different genetic routes, as the constellation of alleles that respond to selection quickest will be somewhat different due to drift among populations (Barton, 1989). Therefore, it may be the case that polygenic traits are even less susceptible to a shared genetic basis to adaptation across populations than simple traits. However, we currently lack good models and methods with which to test this.

### Are species held together by widespread selective sweeps?

Our results touch on an old debate on the evolutionary coherence of species. Mayr and many others have argued that species are coherent evolutionary units because they are united by shared gene flow (pages 521–522 in Mayr, 1963). However, this argument has been questioned by a number of authors based on relatively high levels of differentiation, and low rates of dispersal, in many species (Ehrlich & Raven, 1969; Levin, 1979). Even if gene flow is not high enough to prevent neutral differentiation or local adaptation, a number of authors have argued that species are cohesive if gene flow is high enough for globally selected alleles (and their hitchhiking haplotypes) to spread across entire species (see also Rieseberg & Burke, 2001; Morjan & Rieseberg, 2004; Ellstrand, 2014). At present, large scale genotyping and sequencing projects, along with more sophisticated methods, are highlighting ever more signals of gene flow between populations and species (Patterson et al., 2012; Sousa & Hey, 2013; Hellenthal et al., 2014). However, our work on geographic convergent adaptation (see also Ralph & Coop, 2010, 2014) suggests that species should often adapt to widespread selection pressures through convergent evolution rather than waiting for a single allele to migrate across the range.

In support of this idea, putative recent selective sweeps seem to often be geographically restricted (Pickrell et al., 2009; Coop et al., 2009; Granka et al., 2012), rather than species-wide (but see Clark et al., 2007; Long et al., 2013, for a potential example). This is likely in part due to the relatively low incidence of widespread selection pressures, but as noted above even when we know of widespread selection pressures (e.g. malaria) the response is usually convergent, not shared, across large spatial scales. On the other hand, introgression of adaptive alleles across species and sub-species boundaries, suggests that selected alleles do sometimes spread despite low migration rates (see Hedrick, 2013, for a recent review). However, at least some of these cases may be caused by introgression of haplotype complexes consisting of many, tightly linked, beneficial alleles (that perhaps are inaccessible by mutation over reasonable time-scales for a population in a new environment). Currently we can only scan genomes for species-wide sweeps in those few organisms with population-scale sequence data, and so we do not know if these observations generalize to most species. This is rapidly changing, and will allow us to form a much improved picture of the relationship between the level of neutral population structure, and the age and geographic spread of selected alleles across many species.

Even if selective sweeps only bring alleles to fixation locally, they are still potentially a stronger homogenizing force than neutral mixing through migration. Under neutral mixing, the mean number of generations back to the most recent common ancestor is on order of the total effective population size. This quantity has not been worked out for a model with simultaneous local sweeps, but will be somewhat analogous to the “spatial Λ-Fleming–Viot” models of Barton et al. (2013b), in which local sweeps occur independently across the range. Lineages that are closely linked to the sweeping allele (∼ *v/χ* Morgans) will be moved towards the center of the sweep (a displacement *O*(*χ*)), and pairs of lineages caught up in the same sweep could be forced to coalesce (see Barton et al., 2013a, for work on geographic hitchhiking). In this case lineages and alleles are literally hitchhiking across space. The overall rate of lineage movement and coalescence depends on the rate of sweeps, their geographic scale, and the rate of recombination, and could be calculated by combining the result presented here with Barton et al. (2013b) and Barton et al. (2013a). However, if geographic sweeps are common then this may substantially speed up the rate of mixing compared to neutral drift and migration.

### How then do substitutions occur?

If it is rare for gene flow to rapidly spread selected alleles across a species range, how then do selected alleles ever become fixed within species? Drift alone will act only slowly to sort variants within species into divergence among species. Slight selective differences in the pleiotropic effects (or linked background) among convergent alleles could allow one allele to press into areas occupied by other alleles. Furthermore, repeated bouts of adaptation in particular genomic regions may act to push a subset of previously selected alleles to fixation across the species range, through the spread of the genetic backgrounds on which they arise. However, it seems likely that this is a slow process compared to the initial rapid spread of selected alleles.

### Speciation and extinction as phases of molecular evolution?

One potential resolution is that many selected alleles achieve fixation, not through their own species-wide spread, but rather through subsequent large-scale changes in geographic range size induced by extirpation of the species over parts of its range (see Barton et al., 2013b, and references therein for how such a model could be constructed). Such drops in range size may fix, or radically change the species-wide frequency, of alleles previously restricted to small portion of a species range. Furthermore, many modes of speciation are proposed to occur through a geographically-limited subset of populations forming the basis of new species, e.g. the splitting off of part of the range through a vicariance event or dispersal of a subset of individuals. In this case, speciation will cause geographic assortment of polymorphic ancestral variation, again acting to fix variants within newly formed species that were previously polymorphic across ancestral species ranges.

Such ideas are not completely new and represent a perhaps logical consequence of an allopatric or parapatric view of the biogeography of speciation. However, it is worth revisiting this idea as geographically broad population genomic sampling allows us to return to themes in biogeography. Along similar lines, Futuyma has argued that much of the adaptive differentiation within species, e.g. adaptation to local conditions, may be ephemeral and subject to loss due to local extinction and the mixing following the collapse of population structure (Futuyma, 2010, 1987). Futuyma offered this as an explanation of the pattern of punctuated equilibrium (Eldredge & Gould, 1972), and argued that the observation of stasis and rapid anagenesis associated with speciation were consistent with micro-evolution. Futuyma argued that despite rapid adaptation over short time-scales, we may observe morphological stasis in the fossil record as much of this adaptation is lost to local extinction and the collapse of population structure (see also Lieberman & Dudgeon, 1996; Eldredge et al., 2005; Futuyma, 2010). Furthermore, he suggests that speciation may act as ratchet to prevent the loss of differentiation, acting to maintain adaptive changes among populations, and prevent their loss by inter-breeding. At face value the rate of species formation seems too low to contribute to this process. However, Rosenblum et al. (2012), and many others, have argued that the rate of speciation may well be quite high, but that the majority of incipient species do not persist long due to reabsorption or extinction. Changes in range size, due to local extinction, can also be very rapid on the time-scales over which alleles may spread on the landscape (Gaston, 2003; Hewitt, 1996). Repeated bouts of extinction and speciation will send waves of alleles to fixation along particular lineages.

Such a link between speciation and substitution would not imply that substitutions should necessarily be thought of as being clustered at splits in inferred phylogenies (see Pennell et al., 2014a; Venditti & Pagel, 2014; Pennell et al., 2014b, for a recent exchange on this). Neutral substitutions are unaffected by this process, because they accumulate in a clocklike manner along lineages, as dictated by the mutation rate, regardless of the geographic details of their polymorphic stage. Turning to the accumulation of adaptive substitutions, it is likely that splits in phylogenies are only a tiny proportion of all incipient speciation events, because extinction rates may be high (Rosenblum et al., 2012), and so every lineage has likely passed through many “speciation events” in addition to the observed ones. Under these assumptions, spatial polymorphisms could accumulate gradually in geographically restricted populations between the large-scale biogeographic events that cause their fixation or loss. This effectively decorrelates the time at which new alleles arise and when they fix in the species, an effect similar to that pointed out by Gillespie (1994).

This view would not imply that adaptive evolution or speciation is driven by the shifting balance or genetic revolutions (Wright, 1932; Mayr, 1954), whereby genetic drift allows populations to cross fitness valleys and substitute novel epistatic combinations. Although geographic lineage sorting via speciation and extinction can be thought of as very large-scale genetic drift events, in the models we study here the initial spread of alleles is due to selection, not drift (see also Futuyma, 1989, for discussion).

There is evidence that a reasonable fraction of genome-wide substitutions are fixed by positive selection in a number of species (most notably Drosophila, Sella et al., 2009). Under the geographic view of fixation, selection has played a strong role in the establishment of these alleles locally. As we get more broadly geographic population genomics sampling for a range of species we will have the opportunity to study whether the class of alleles that contribute to local differentiation are similar to those underlying species divergence, and the extent to which the answer to this depends on the age and type of population structure within species.

Finally, we close by noting that range expansion and speciation are obviously not separate from adaptive differentiation. The invasion of new geographic areas may lead to a burst of adaptive differentiation, at least in a subset of genes, and speciation may be associated with rapidly adaption to novel environments. Conversely, if the geographic spread of adaptive alleles within ranges is slow (e.g. if they only offer a local advantage or if they are selectively excluded) this may allow Dobzhansky-Muller incompatibilities to arise within species, effectively offering a mechanism for hybrid incompatibilities to evolve in parapatry, and fracturing the range (Bank et al., 2012; Kondrashov, 2003; Bierne et al., 2011). The alleles that act as components in many of the Dobzhansky-Muller incompatibilities studied to date are geographically restricted (see Cutter, 2012). Therefore, it seems possible that populations within species may often be tending towards speciation, and that as outlined here that this may drive some proportion of molecular divergence.

## Acknowledgements

We thank Michael Turelli, Jon Seger, and the Coop lab for helpful conversations. We thank Florence Débarre, Douglas Futuyma, Michael Whitlock, Molly Przeworski, and Sam Yeaman for helpful comments on an earlier version of the paper. All figures and calculations were performed in R (R Core Team, 2014). The figure color paletes were generated using Karthik Ram’s “Wes Anderson” R color palette https://github.com/karthik/wesanderson. This work was supported by grants from the National Science Foundation under Grant No. 1262645 to PLR and GC and by the National Institute of General Medical Sciences of the National Institutes of Health under award numbers NIH RO1GM83098 and RO1GM107374 to GC.

